# The genetic architecture of maize yellow mosaic virus resistance in corn

**DOI:** 10.64898/2026.07.17.739229

**Authors:** Erik W. Ohlson, Christopher A. Nacci, Nitika Khatri, Kristen J. Willie, Jennifer R. Wilson

## Abstract

Maize yellow mosaic virus (MaYMV) is an emerging polerovirus of corn and other grass species. Due to its broad host range and transmission by multiple aphid species, management strategies such as crop rotations, pesticides, and weed control are likely ineffective. Therefore, resistant cultivars are needed. In this study, we characterized the Goodman 282 maize diversity panel for its response to MaYMV. Leaf reddening symptoms were quantified, and diagnostics were performed to assess infection and obtain a semi-quantitative measure of virus titer. Low titer lines and inbreds representing symptomatic and asymptomatic infected phenotypic classes were characterized further by RT-qPCR. Genome-wide association studies were performed using MLM, FarmCPU, and BLINK models to identify SNPs associated with disease. In total, 64% of lines were asymptomatically infected. Although all lines tested positive for infection by MaYMV in at least one experiment, Ky226 had reduced viral titer compared to other lines. Sixteen quantitative trait nucleotides (QTN) were identified, many of which are linked to genes implicated in flavonoid, carotenoid, and phenolic metabolic pathways as well as antiviral defense. Notably, a QTN associated with a chalcone synthase, a key enzyme in the flavonoid biosynthesis pathway, was detected by even the most conservative, MLM model. These results indicate that the genetic architecture of corn in response to MaYMV is complex, and that developing immune cultivars may not be achievable using natural sources of genetic variation. However, this study provides a foundation for breeding maize with improved tolerance and advances our understanding of host response to MaYMV infection.

## Introduction

Maize yellow mosaic virus (MaYMV) is an emerging corn virus first identified in the Yunnan, Guizhou, and Anhui provinces of China in 2013 (Chen et al. 2016; Wang et al. 2016). Since its discovery, MaYMV has been detected across Asia (Lim et al. 2018b; Nithya et al. 2021; Chen et al. 2026), Sub-Saharan Africa (Guadie et al. 2017; Yahaya et al. 2019; Palanga et al. 2017; Read et al. 2019; Wamaitha et al. 2018; Welgemoed et al. 2020; Adams et al. 2017; Massawe et al. 2018), and both North (Lappe et al. 2022) and South America (Bernreiter et al. 2017; Gonçalves et al. 2017). MaYMV is especially prevalent in tropical and subtropical regions, where its aphid vectors and alternative plant hosts are abundant. In Sub-Saharan Africa, MaYMV was detected in 57-78% of plants tested with virus-like symptoms (Adams et al. 2017; Massawe et al. 2018; Tanui et al. 2021; Guadie et al. 2017; Asiimwe et al. 2020), and in China up to 52% of sugarcane samples with mosaic symptoms tested positive for MaYMV (Sun et al. 2021).

MaYMV’s widespread distribution and prevalence is facilitated by its broad host range and the numerous aphid species with transmission capacity. Although it was first detected in corn (Chen et al. 2016; Wang et al. 2016), it also infects economically important species including sugarcane (*Saccharum* spp.) (Sun et al. 2021; Yahaya et al. 2017), sorghum (*Sorghum bicolor*) (Lim et al. 2018b; Wamaitha et al. 2018), wheat (*Triticum aestivum*) (Guo et al. 2022), barley, oats, rye (Ohlson et al. 2024), and millet (*Panicum miliaceum*) (Lim et al. 2018b) as well as many other weedy hosts in the *Poaceae* family (Yahaya et al. 2017; Ohlson et al. 2024). MaYMV may be spread among and between these various hosts by aphid species including *Rhopalosiphum maidis* (Gonçalves et al. 2020), *R. padi* (Stewart et al. 2020), and *Schizaphis graminum* (Ohlson et al. 2024) in a circulative non-propagative manner (Sun et al. 2021). The breadth of hosts and vectors involved in the epidemiology of MaYMV highlights the difficulty of controlling the disease using traditional pest management strategies such as crop rotations and pesticides.

MaYMV is a yellow dwarf virus, along with the economically important maize, wheat, and cereal yellow dwarf viruses in genus *Polerovirus* and barley yellow dwarf viruses in the genus *Luteovirus.* Symptoms of MaYMV infection are similar to those induced by these other yellow dwarf viruses, with leaf tip reddening and yellow mosaic symptoms most frequently reported, followed by stunting, chlorosis, vein clearing, leaf curling, and an overall reduction in plant health (Gonçalves et al. 2017; Lim et al. 2018a; Wang et al. 2016; Shi et al. 2022; Stewart et al. 2020; Stewart and Willie 2021; Sun et al. 2019). Although the economic impact of MaYMV is poorly characterized, in controlled growth chamber studies, MaYMV reduces plant vigor by 10% (Stewart and Willie 2021). MaYMV is also commonly found in co-infections with other economically significant viruses of maize including maize chlorotic mottle virus, maize rayado fino virus, and sugarcane mosaic virus (SCMV) (Sun et al. 2019; Tanui et al. 2021; Yahaya et al. 2017; Chen et al. 2016; Gonçalves et al. 2017), where yield losses of 10 - 30% were estimated (Bernreiter et al. 2017). Given the widespread distribution of MaYMV and its potential impact on plant health, management strategies are desirable to address this emerging disease.

Since conventional spray and rotation strategies are ineffective at controlling the spread of MaYMV due to its broad host range and effective transmission by several aphid vectors, one of the most promising strategies for mitigating its impact is by breeding and deploying resistant or tolerant maize cultivars. Rapid development of resistant cultivars is facilitated by molecular breeding technologies including gene editing or conventional marker assisted breeding, both of which require the identification of the causal resistance genes or associated genomic loci. Therefore, in this study we evaluated the Goodman 282 maize diversity panel for its response to MaYMV and performed a genome wide association study (GWAS) to identify potential genes and loci of interest, providing maize breeders with new tools to combat this emerging maize virus disease.

## Materials and methods

### Plant, pathogen, and insect materials

The Goodman 282 diversity panel, which captures a large proportion of the genetic diversity present among public sector maize breeding programs worldwide, was evaluated for its response to MaYMV (Flint-Garcia et al. 2005; Gerdes and Tracy 1993). The population consists of stiff stalk, non-stiff stalk, tropical, popcorn, and mixed origin inbred lines (Table S1). Among the 277 lines in the population, 18 were not included in this study due to poor germination or difficulty regenerating seed under temperate field conditions. Seeds were originally obtained from the North Central Regional Plant Introduction Station (Ames, IA, USA) and each line has been maintained by single-seed descent. Oh28, an MaYMV susceptible line with strong leaf reddening symptoms was included in all experiments as a susceptible control.

A single MaYMV isolate was used for all experiments in this study. The isolate was originally collected from Morogoro, Tanzania and has been maintained in susceptible corn by serial transmission using *R. maidis* or by vascular puncture inoculation of seed with cryo-preserved MaYMV infected leaf tissue (Asiimwe et al. 2020; Stewart et al. 2020). The MaYMV isolate used in this study was previously shown to induce severe leaf reddening symptoms in some susceptible corn lines and is aphid-transmissible at a high rate (Stewart et al. 2020; Ohlson et al. 2024). However, no mosaic, chlorosis, or other visible symptom types are associated with infection by this isolate in any of the lines tested previously in our experimental conditions.

The *R. maidis* colony used for all experiments was obtained from Dr. Anna Whitfield at the North Carolina State University and was originally collected near Manhattan, Kansas, USA. Inoculative aphids for each experiment were obtained by first taking 5 - 10 aphids reared on MaYMV infected source plants, inoculating 9 - 11-d old susceptible Oh28 maize plants, and then caging those plants in BugDorms. After a 6 - 8-d inoculation access period (IAP), these plants were fumigated with Nuvan ProStrips (18.6% dichlorvos) in large plastic bins for 3.5 - 4-h to destroy live aphids. The inoculated plants were then grown for 13-d for disease to develop. Following this incubation period, leaf clippings containing 30 - 40 nonviruliferous aphids were placed in the whorl of each MaYMV infected plant and given a 20 - 22-d acquisition access period (AAP). Aphids from these plants were subsequently used for experimental inoculations.

### Evaluation of disease response in the Goodman 282 diversity panel

Disease response among lines in the diversity panel was evaluated across multiple experiments. All experiments were conducted under controlled growth chamber conditions, with day-night temperature setpoints of 25 °C and 21 °C respectively, a 14:10-hr day:night period, and light intensity setting of 600 µmol/m^2^/s.

To obtain detailed symptom data for each line across multiple time points, plants were evaluated in 10-cm pots divided across 10 experiments due to the intensive labor requirements associated with inoculations and disease evaluations. Five seeds were planted in each pot in two replicates. An uninoculated pot was also planted for each line to discern potential differences in response to disease compared to the diverse physiological characteristics present within the population. Plants were inoculated at the V2-V3 growth stage, 9 - 10-d after planting, with five viruliferous aphids applied to the whorl of each individual plant. The plants were caged in BugDorms for a 4 - 5-d IAP, after which the plants were fumigated with Nuvan ProStrips and cages were removed. Pots were sprinkled with 1% granular marathon (OHP) to ensure no live aphids persisted. Symptom data were collected for each experiment three times between 14 - 31-d post inoculation (dpi) and leaf tissue was collected following the final evaluation for diagnostics. A quantitative measurement of leaf tip reddening area was taken at each time point by measuring reddening length in centimeters for each symptomatic leaf. Using this data, an area under disease progress curve (AUDPC) was calculated for each plant and the average AUDPC was calculated for each line as:

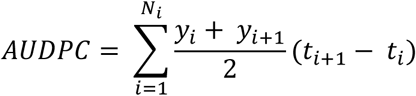

To confirm infection by MaYMV, tissue samples were pooled from each pot and ground in grape extraction buffer (1:20 w/v; GrEB: 0.05 M sodium carbonate buffer pH 9.6, 2% polyvinylpyrrolidone-40, 0.2% bovine serum albumin, 0.05% Tween-20) then 4 ul of extract was denatured in GES buffer (0.1 M glycine-NaOH pH 0.0, 50 mM NaCL, 1 mM EDTA, 0.5% Triton X-100, and 10 mM DTT) for 10-min at 95 °C. A one-step RT-PCR was performed as described previously (Xie et al. 2020) to detect the presence of MaYMV using forward primer DM03: 1944F (5’- GGGAGGTTACTTGCAGAGGG-3’) and reverse primer DM04: 2836R (5’-ATGCACTTCAACCAACACGC-3’) primer sequences (Massawe et al. 2018). The thermocycler protocol consisted of a 45-min incubation period at 52 °C for reverse transcription, denaturation at 94 °C for 2-min, followed by 35 cycles of 15-s denaturation (94 °C), 30-s annealing (60 °C), and 60-s extension (72 °C) with a final extension of 7-min (72 °C). The PCR product was visualized on a 1% agarose gel containing 0.25 μg/mL ethidium bromide by UV light using a gel imaging system to detect the presence or absence of an MaYMV amplicon.

To capture quantitative variance of the disease response and virus titer present among the entire diversity panel simultaneously within individual experiments, the entire population was planted in two replicated experiments in 5-cm Deepots (Stuewe & Sons, Inc). Two seeds per line were planted in each Deepot and thinned to a single plant prior to inoculation with MaYMV at V2-V3 growth stage 9-d after planting. Five aphids were transferred to the whorl of each plant and covered by individual plastic tube cages to prevent aphids from escaping. After a 5-d IAP, the plants were fumigated with Nuvan ProStrips, the tube cages were removed, and the soil was sprinkled with Marathon. At 27 dpi, plants were evaluated for disease symptoms using the previously described method. Tissue was collected 28 dpi to semi-quantitatively assess virus titer using a protein A antibody sandwich enzyme-linked immunosorbent assay (PAS-ELISA). A total of 20 6.35 mm leaf punches were collected across the top 4 - 5 leaves of each plant. Leaf punches were evenly distributed across each leaf to obtain a representative sampling. The leaf disks were collected in 2 mL microtubes with six 2-mm steel beads in each tube, flash frozen in liquid nitrogen, and stored at -80 C prior to processing the samples. Prior to PAS-ELISA, the samples were cryo-ground using a SPEX SamplePrep 1600 MiniG tissue homogenizer for 1-min. PAS-ELISA was conducted as previously described (Wilson et al. 2025). Plates were coated overnight at 4°C with Protein A (1 ug/mL) in carbonate coating buffer (CCB, Agdia Inc.). After washing with phosphate-buffered saline plus 0.05% Tween 20 (PBS-T), plates were incubated with the cross adsorbed anti-MaYMV antibody developed in Wilson et al. (2025) (Wilson et al. 2025) diluted 1:1000 in PBS-T plus 3% bovine serum albumin (BSA). Cryoground samples were re-suspended in 600 uL General Extract Buffer (GEB, Agdia Inc.) and after plates were again washed with PBS-T, 100 uL of extract was applied to the plate in technical duplicate and incubated overnight at 4°C. After washing with PBS-T, the anti-MaYMV antibody was applied again, diluted 1:1000 in PBS-T with 3% BSA. Next, after washing, Protein A conjugated to alkaline phosphatase was applied at a dilution of 1:5000 in PBS-T plus 3% BSA. After the final wash, PNPP substrate dissolved in PNP substrate buffer (Agdia Inc.) was applied and the colorimetric reaction was monitored over several hours, taking absorbance readings at 405 nm using a Synergy H1 microplate reader (BioTek). Samples were considered positive for MaYMV if the absorbance reading was more than twice that of the uninfected controls. To account for variability between plates, the mean absorbance of each line was normalized by dividing by the mean absorbance of the corresponding Oh28 susceptible controls that were included on each plate.

### PAS-ELISA and RT-qPCR comparisons of resistant and susceptible lines

Three lines, CML103, CML333, and Ky226, that tested negative by ELISA based on an absorbance threshold of 2x the uninfected control absorbance were selected for further testing. Four susceptible lines were also evaluated including the symptomatic susceptible lines CML333 and control Oh28, as well as B57 and Mo18W, two non-symptomatic lines that were definitively positive by ELISA. Each of these lines were planted in Deepots in five replicates and inoculated with MaYMV as previously described. At 28 dpi, individual plants were sampled separately for RT-qPCR and ELISA testing and subsequently cryo-ground as described above.

PAS-ELISA was conducted on the samples as described earlier. RNA was isolated from the second sample per plant using the Direct-zol RNA Miniprep Kit (Zymo Research) following manufacturer’s instructions with on-column DNaseI treatment. Complementary DNA was synthesized using iScript Reverse Transcription Supermix for RT-qPCR (Bio-Rad) using 0.4 ug of RNA and following the manufacturer’s instructions. Quantitative PCR was conducted using 5 uL of a 1:5 dilution of cDNA into SsoAdvanced Universal SYBR Green Supermix (Bio-Rad) using the primers MaYMV 3348F (5’- GTGAGTTGCAAGTGCTGGAA - 3’) and MaYMV 3546R (5’- TCCTAGCTCTGCGTCCATTT - 3’). MaYMV titer is expressed relative to Oh28 with normalization to two reference genes, *UBCP* and *FPGS* (Manoli et al. 2012) using the method described by Vandesompele *et al*. 2002 (Vandesompele et al. 2002). For each assay, data were evaluated for normality using the Shapiro-Wilk test, and homogeneity of variance was assessed using Levene’s test. Based on these results, RT-qPCR data were analyzed using Welch’s ANOVA followed by Games-Howell *post-hoc* comparisons as the data were non-normal and heteroscedastic. The ELISA data were analyzed using one-way ANOVA followed by Tukey HSD since they met the assumptions of normality and homoscedasticity.

### Genome Wide Association Studies

Unimputed SNP variants, called from the publicly available 7x whole genome sequencing data for each line in the 282 panel were downloaded from CyVerse data commons (/iplant/home/shared/panzea/hapmap3/hmp321/unimputed/uplifted_APGv4) (Bukowski et al. 2017). The SNPs were uplifted to version five of the B73 reference genome (Zm-B73-REFERENCE-NAM-5.0) using CrossMap (Zhao et al. 2014) and the v4 to v5 chain file obtained from maizeGDB.org. Variants were filtered to include only biallelic SNPs with call rates ≥ 0.8, minor allele frequency (MAF) ≥ 0.05, and the local linkage disequilibrium (LLD) flag. Variants flagged with the near indels within 5 bp (NI5) annotation were excluded from analysis and all heterozygous SNP calls were set to missing. SNP imputation was performed using Beagle 5.5 (Browning et al. 2018) with 10 burn-in iterations, 15 genotype phase iterations, an effective population size (N_e_) of 50,000, and using the NAM high resolution genetic maps obtained from maizeGDB.org to estimate the genetic distances between physical genome coordinates (Yu et al. 2008; McMullen et al. 2009). Only biallelic variants with MAF ≥ 0.05 in the population were included in the final analysis. Pruning was conducted using PLINK version 2.0 to include only SNPs with linkage disequilibrium (r^2^) < 0.99 for the GWAS analyses and an r^2^ < 0.1 was used to calculate kinship and structure. In total, 5,245,123 SNPs were retained for GWAS analysis, and 181,580 of these SNPs were used to calculate principal components and kinship in GAPIT Version 3 using the default parameters (Wang and Zhang 2021).

GWAS was conducted in GAPIT using three models: mixed linear model (MLM), FarmCPU, and BLINK (Wang and Zhang 2021) to identify SNPs significantly associated with AUDPC (10-cm pot experiments), disease severity (DS) (Deepot experiments), and the normalized ELISA values. The first three principal components were used as covariates as determined by scree test and the kinship matrix was provided for MLM to account for relatedness between lines. A stringent Bonferroni corrected logarithm of the odds (LOD) threshold of 8.02 (p ≤ 0.05) was used to identify significant SNPs with a suggestive threshold delineated at -log_10_(1/5,245,123). Annotations were extracted from the version 5 of the B73 reference genome within 50 kb on either side of significant quantitative trait nucleotides (QTN) and extended to 75 kb if no gene annotations were present as was the case with a single QTN.

## Results

### Disease response to MaYMV in the Goodman 282 maize diversity panel

In the 10-cm pot experiments, MaYMV reddening symptoms were detected in 65 of the 258 lines that were evaluated. The average AUDPC was 36.4 ± 25.1 among the symptomatic lines indicating significant variation in disease response. Similarly, 53 of the 250 lines evaluated in the Deepot experiments displayed MaYMV symptoms, with an average DS of 9.3 ± 8.1 (Fig. 1). Cumulatively, 167 of the 259 unique lines tested in this study were free of the reddening symptoms associated with MaYMV despite evidence of virus infection based on RT-PCR and ELISA. No other symptoms were found to be associated with infection by the MaYMV isolate used in these experiments regardless of genetic background.

**Fig. 1.**
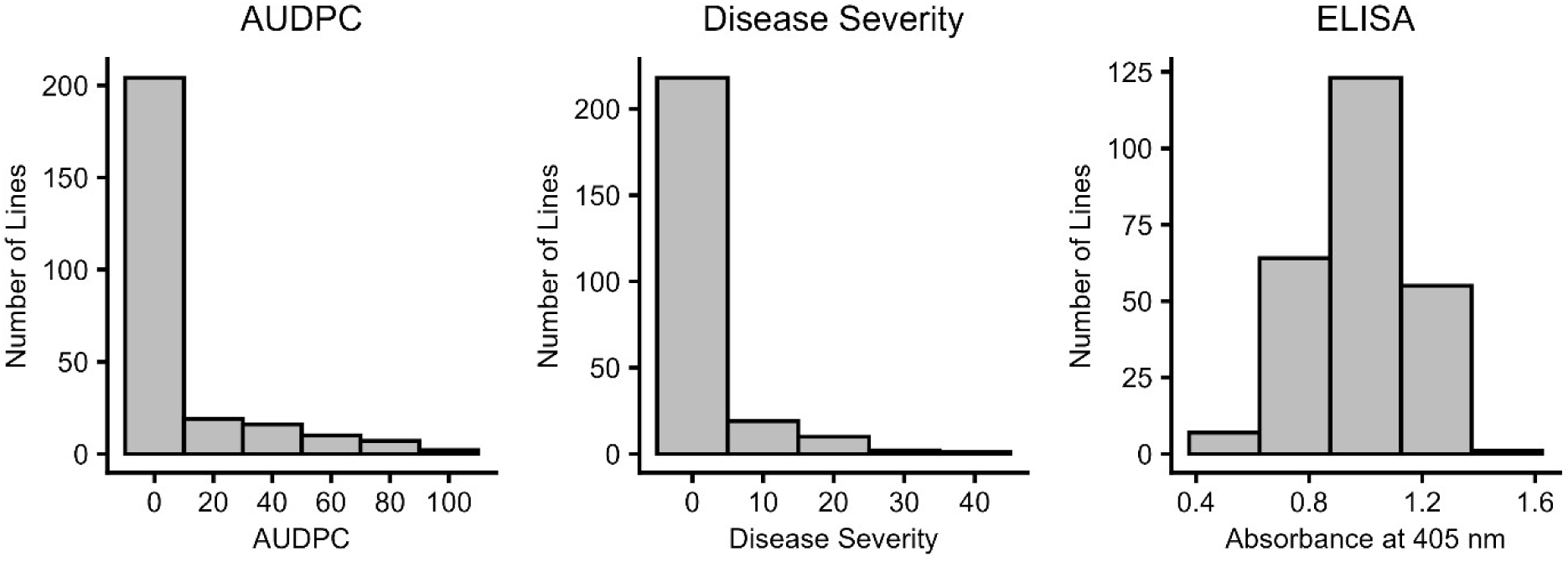
Phenotypic distribution of response to inoculation with maize yellow mosaic virus in the maize 282 association mapping panel. Histograms show the distributions for area under disease progress curves (AUDPC) and final disease severities based on leaf reddening response as well as measures of sample absorbance at 405 nm as tested by enzyme-linked immunosorbent assay (ELISA).

Both RT-PCR and PAS-ELISA diagnostics indicated that none of the lines evaluated in this study are immune to MaYMV infection, with all lines testing positive by RT-PCR even if they were negative by ELISA. The average relative ELISA absorbance in the population when normalized to the Oh28 highly susceptible control line, was 0.98 ± 0.17 (Fig. 1). Furthermore, average ELISA absorbances were similar among symptomatic and non-symptomatic lines, averaging 0.99 ± 0.17 and 0.98 ± 0.18 respectively in each phenotypic class. Although they tested positive by RT-PCR, CML45, CML103, Ky226, and CML287 were not initially considered positive by ELISA in either of the two experimental replicates conducted in Deepots based on the 2x the healthy control absorbance threshold, warranting further investigation as a potential source of MaYMV resistance.

### Genome wide association studies

GWAS analyses detected several SNPs that were highly associated with reddening responses by measures of AUDPC and final DS. While the highly conservative MLM method detected the presence of only a single QTN exceeding the Bonferroni corrected threshold on chromosome 6 for AUDPC, suggestive QTN on chromosome 4 for AUDPC, and chromosomes 5, 7, and 10 for disease severity were identified, for each of which significant associations were detected by the more powerful FarmCPU and BLINK models (Table 1, Fig. 2). This chromosome 6 QTN was tightly linked to five genes including a 3-ketoacyl-CoA synthase gene and explained 63% of the phenotypic variance (PVE) in the MLM analysis and 20% in FarmCPU (Table S2).

**Fig. 2.**
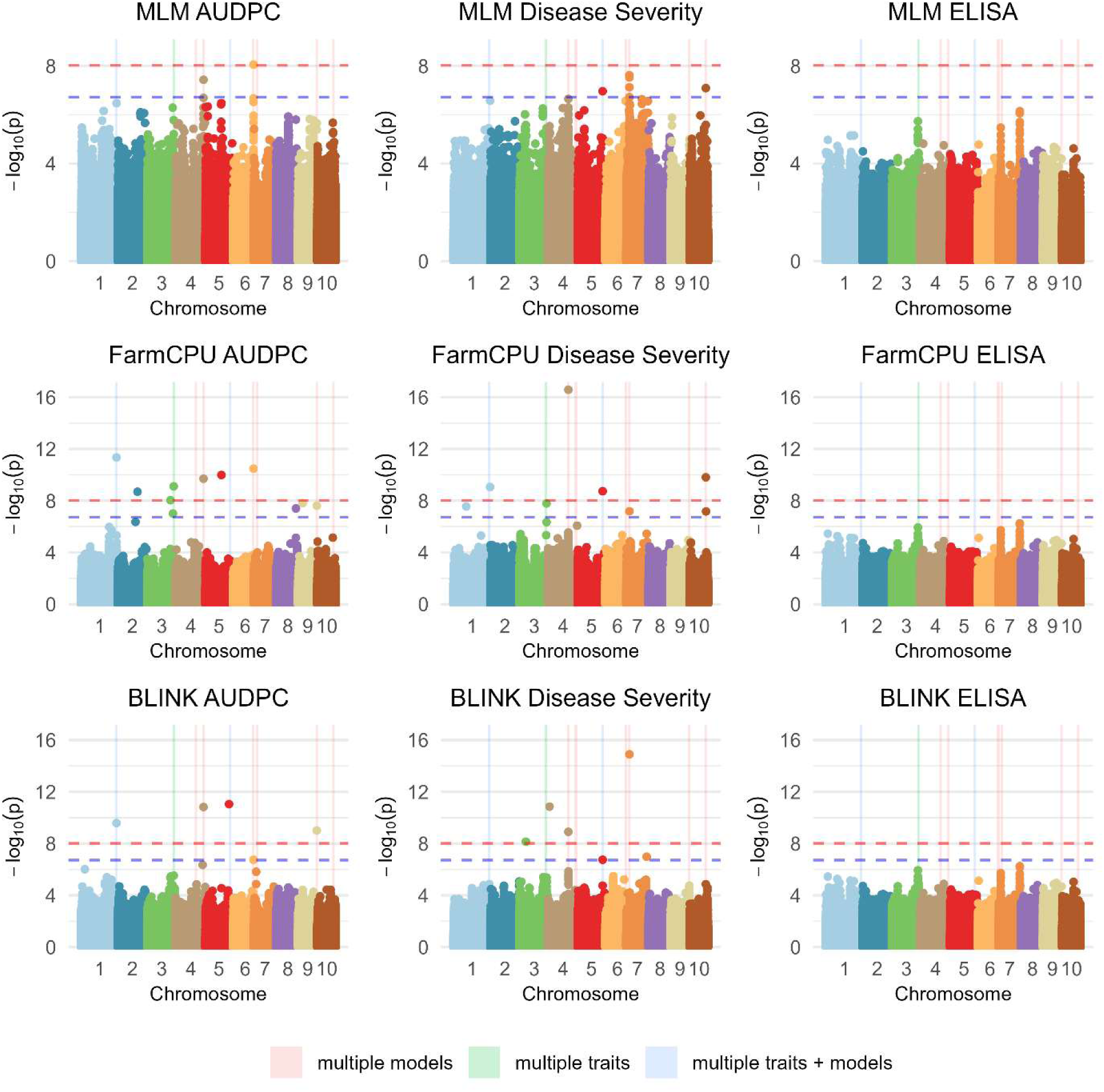
Manhattan plots for single nucleotide polymorphisms (SNP) associated with response to maize yellow mosaic virus inoculation in the maize 282 association mapping panel. Associations were tested between markers and area under disease progress curves (AUDPC), disease severity, and enzyme linked immunoassay absorbance (ELISA) using three models: mixed linear models (MLM), Fixed and random model circulating probability unification (FarmCPU), and Bayesian-information and Linkage-disequilibrium Iteratively Nested Keyway (BLINK) and significant loci across multiple models and/or traits are highlighted. Bonferroni corrected significance [-log10(0.05/5,245,123] and suggestive thresholds [- log_10_(1/5,245,123)] are indicated by red and blue horizontal lines respectively.

**Table 1.** Single nucleotide polymorphisms (SNPs) and genes associated with maize yellow mosaic virus resistance.

| Chr | Position | Trait | Model | p-value | Effect | PVE | MAF | Gene Annotation | Description |
| --- | --- | --- | --- | --- | --- | --- | --- | --- | --- |
| 1 | 295,708,934 | AUDPC | BLINK | 2.6E-10 | -11.2 | 20.8 | 0.10 | Zm00001eb061060 | H15 domain-containing protein; Histone H1 |
|  |  |  | FarmCPU | 4.5E-12 | -9.8 | 17.6 |  |  |  |
| 1 | 305,881,404 | DS | FarmCPU | 8.8E-10 | -1.8 | 14.7 | 0.09 | Zm00001eb064730 | Nuclear factor Y subunit C10; Transcription factor CBF/NF-Y/archaeal histone domain-containing protein |
| 2 | 162,879,044 | AUDPC | FarmCPU | 2.0E-09 | 4.7 | 2.6 | 0.16 | Zm00001eb095000 | Cytochrome b5 heme-binding domain-containing protein |
| 3 | 53,304,527 | DS | BLINK | 7.1E-09 | -2.3 | 17.3 | 0.08 | Zm00001eb130080 | ATP citrate synthase |
| 3 | 195,517,056 | AUDPC | FarmCPU | 9.1E-09 | -5.3 | 7.6 | 0.10 | Zm00001eb151300 | Receptor-like serine/threonine-protein kinase |
| 3 | 222,494,308 | AUDPC | FarmCPU | 7.8E-10 | -10.1 | 20.1 | 0.04 | Zm00001eb159250 | Cytochrome P450 |
| 4 | 13,865,340 | DS | BLINK | 1.4E-11 | -2.2 | 16.7 | 0.11 | Zm00001eb168440 | Multiple organellar RNA editing factor 1 mitochondrial; DAG protein; Uncharacterized protein |
| 4 | 171,091,807 | DS | BLINK | 1.2E-09 | -1.9 | 14.6 | 0.15 | Zm00001eb189970 | Fcf2 pre-rRNA processing protein |
|  |  |  | FarmCPU | 2.7E-17 | -2.4 | 15.9 |  |  |  |
| 4 | 234,458,912 | AUDPC | BLINK | 1.5E-11 | -10.6 | 35.7 | 0.07 | Zm00001eb204380 | Uncharacterized protein |
|  |  |  | FarmCPU | 2.0E-10 | -8.6 | 11.2 | 0.07 |  |  |
| 5 | 133,996,275 | AUDPC | FarmCPU | 1.0E-10 | -7.1 | 6.1 | 0.09 | Zm00001eb237110 | Protein DETOXIFICATION (Multidrug and toxic compound extrusion protein) |
| 5 | 198,022,994 | AUDPC | BLINK | 8.9E-12 | -6.6 | 6.3 | 0.21 | Zm00001eb250080 | Geranylgeranyl pyrophosphate synthase4 |
| 5 | 207,921,461 | DS | FarmCPU | 1.9E-09 | -2.0 | 17.0 | 0.07 | Zm00001eb251670 | MYB-CC type transcription factor LHEQLE-containing domain-containing protein; G2-like transcription factor (Homeodomain-like superfamily protein) |
| 6 | 176,624,116 | AUDPC | FarmCPU | 3.3E-11 | -9.3 | 20.3 | 0.05 | Zm00001eb296200 | Major facilitator superfamily profile domain-containing protein; Solute carrier family 2, facilitated glucose transporter member 8 (Sugar transporter ERD6-like) |
|  |  |  | MLM | 8.8E-09 | -17.6 | 63.0 |  |  |  |
| 7 | 25,315,899 | DS | BLINK | 1.3E-15 | -3.0 | 25.0 | 0.10 | Zm00001eb304290 | Calreticulin |
| 7 | 176,333,555 | ELISA/DS | Suggestive | 7.1E-07 | 0.1 | NA | 0.10 | Zm00001eb328050 | E3 ubiquitin-protein ligase RKP |
| 9 | 158,111,554 | AUDPC | BLINK | 9.7E-10 | -5.5 | 12.0 | 0.20 | Zm00001eb402630 | glutathione transferase |
| 10 | 134,445,239 | DS | FarmCPU | 1.6E-10 | -2.6 | 28.4 | 0.06 | Zm00001eb426920 | Metal-nicotianamine transporter YSL7 |

Chr, chromosome; Position, B73 ref v5 position of SNP in base pairs; DS, Disease severity; AUDPC, Area under disease progress curve; PVE, % phenotypic variance explained; MAF, minor allele frequency; Gene, closest annotated gene to SNP

FarmCPU detected the highest number of QTN, 11 in total: seven for AUPDC and four associated with DS. Two of the largest effect QTN explained > 15% of the phenotypic variance each for AUDPC and localized to chromosomes 1 and 3, in addition to the aforementioned chromosome 6. Between 2 - 5 genes were present within 50 kb of each of these loci. All four QTN detected by FarmCPU for DS explained between 15 - 28% of phenotypic variance each and were tightly linked to 4 - 9 annotated genes, notably a nuclear factor Y (NFY) on chromosome 1, an aspartyl protease gene on chromosome 4, and a G2-like transcription factor as well as a ubiquinol oxidase on chromosome 9 (Tables 1, S2, Fig. 2).

The BLINK models detected the presence of 8 QTN associated with MaYMV disease response, four for each of AUDPC and DS. Two QTN accounted for 20 - 35% of the phenotypic variance each, corresponding with the same QTN on chromosomes 1 and 4 that were detected by FarmCPU for AUDPC. Each of the DS QTN detected by the BLINK model accounted for between 15 - 25% of the phenotypic variance, one of which colocalized with a FarmCPU QTN for DS at approximately 171 Mbp on chromosome 4, while the other three were located on chromosomes 3, 4, and 7. Between 2 - 3 genes were annotated within 50 kb for each of the unique chromosome 4 and 7 QTN, including a protein containing an eIF-2B subunit on chromosome 4 and a calreticulin gene on chromosome 7. No genes are annotated within 50 kb of the chromosome 3 QTN. However, 53 kb downstream of the significant SNP is an ATP citrate synthase gene (Table 1, S2, Fig. 2).

Although no associations exceeded the Bonferroni or even the suggestive threshold for ELISA absorbance, several obvious LOD peaks were present including two that colocalized with significant or suggestive SNPs for AUDPC and DS near the bottom of chromosome 3 and the top of chromosome 7. The most significant SNP for ELISA by MLM (LOD = 6.1) was located towards the bottom of chromosome 7 and is tightly linked to an E3 ubiquitin-protein ligase. This SNP also exceeded the suggestive value for disease severity across multiple models (Table 1, Fig. 2).

### Comparison of PAS-ELISA and RT-qPCR techniques for quantifying MaYMV titer

Three asymptomatic lines, CML45, CML103, and Ky226, which were identified as potentially resistant (ELISA absorbance < 2x the healthy control) across the two Deepot screening experiments were selected for further study along with the symptomatic susceptible CML333 and control line Oh28. Two ELISA positive but asymptomatic lines, B57 and Mo18W, were also included to explore whether leaf reddening and viral titer were independent from each other. CML287 was not included in this experiment due to insufficient seed viability despite its negative ELISA tests. No significant differences were detected among ELISA absorbance values for any lines in this experiment based on one-way ANOVA (p = 0.39). However, significant differences were detected for RT-qPCR based on Welch’s ANOVA (p = 0.007). Subsequent multiple comparison Games-Howell tests indicated that Ky226 viral titer was significantly lower relative to the symptomatic susceptible control line Oh28 and the non-symptomatic but susceptible B57 (p < 0.05). No other significant titer differences were detected between lines, despite clear variation in symptoms. Only two infected plant samples were obtained for CML45, but despite their lower relative ELISA absorbance, the RT-qPCR relative titers resembled the susceptible controls. Lower ELISA absorbance values were not reproducible in this titer experiment for CML103 compared to susceptible controls, however it displayed more intermediate levels of viral titer by RT-qPCR as it was neither significantly different from Ky226 nor the susceptible Oh28 control line (Fig. 3).

**Fig. 3.**
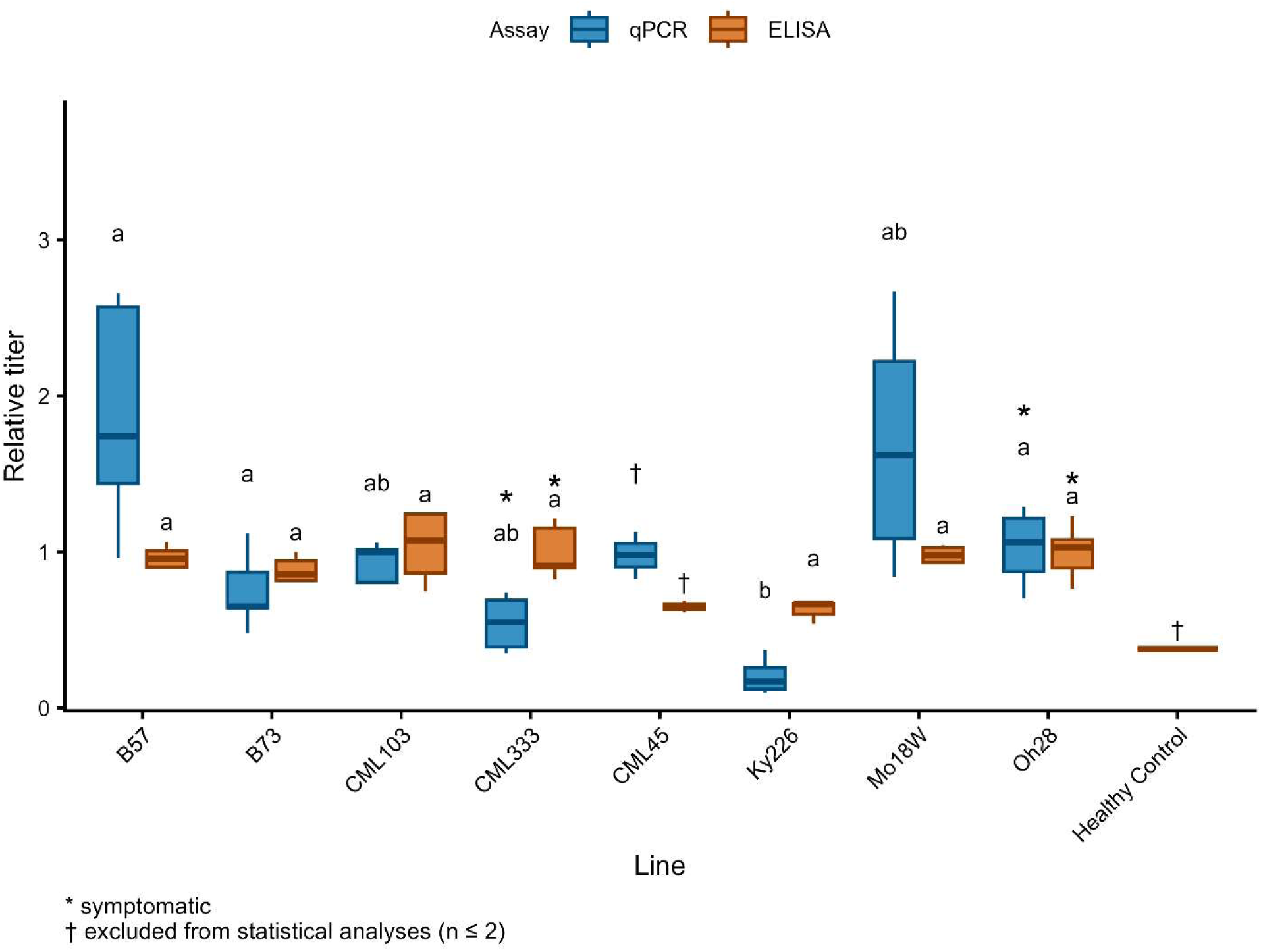
Relative MaYMV titer quantified by two methods in select maize lines. RT-qPCR and PAS-ELISA were conducted to assess MaYMV abundance. RT-qPCR was normalized by corn endogenous control genes FPGS and UBCP. RT-qPCR and ELISA data were analyzed independently. Lines not sharing the same letter within an assay were considered significantly different at alpha = 0.05 Lines not sharing the same letter within an assay were considered significantly different at alpha = 0.05.

## Discussion

MaYMV was found to infect all inbred lines across the entire association mapping population based on ELISA and RT-PCR testing, despite 65% of the lines tested never showing symptoms in any experiment. However, disease severity and, to a lesser extent, viral titer varied significantly across the population. Despite its namesake, the MaYMV isolate used in this study never produced yellowing or mosaic in any of the lines tested. This suggests that previous reports of yellowing and mosaic may be caused by co-infection with another virus such as SCMV, though it is possible that isolate-specific differences or environmental conditions affect MaYMV disease expression. The genetic abundance of asymptomatic lines suggests that developing hybrid varieties without a reddening response may be a straightforward endeavor and could already be the predominant phenotype in hybrid cultivars. However, further study is needed to develop appropriate breeding strategies and determine whether less easily discernable phenotypes under controlled growth chamber conditions, such as reduced yields or grain quality, are associated with asymptomatic infection in single or mixed infections with other pathogens.

In total, we identified 16 unique MaYMV QTN spread across 9 of the 10 maize chromosomes. Several genes within 50 kb of these loci are of particular interest, given their known or predicted functions related to plant defense, disease and stress response, or the biosynthesis of plant phenolics and carotenoids. The most conservative model used in this study, MLM, detected the presence of just a single locus associated with response to MaYMV based on AUDPC on chromosome 6 which explained more than 60% of the phenotypic variance. The annotated genes most tightly associated with this QTN are several major facilitator superfamily (MFS) profile domain proteins, an uncharacterized protein, and 3-ketoacyl-CoA synthase 16 (*kcs16;* Zm00001eb296230), which is one of the most intriguing candidate genes identified in this study. Located ∼24 kb downstream of the significant SNP, *kcs16* is a member of the maize chalcone synthase (CHS) family (Wang et al. 2025). Chalcone synthases are involved in the production of chalcones, organic compounds that serve an important role in plant defense. The catalysis of chalcone synthases serves as the gateway to the biosynthesis of flavonoids, phytoalexins, and other metabolites involved in plant stress response, production of which can cause leaf reddening. It was shown previously that leaf reddening symptoms associated with infection by MaYMV increased expression of *colorless2* (Zm00001eb198030), a chalcone/stilbene synthase N-terminal domain-containing protein (Mlotshwa et al. 2021). This same locus on chromosome 6 was similarly detected for AUDPC by FarmCPU and approached the suggestive threshold for DS by MLM, further supporting the importance of this locus for MaYMV-induced leaf reddening.

While none of the other SNPs identified in this study are as tightly linked with CHS genes as the chromosome 6 locus, several other AUDPC or DS QTN are tightly linked to genes involved in flavonoid, carotenoid, and other pigment biosynthesis pathways. On chromosome 3, BLINK detected a significant DS QTN, most closely linked to the ATP citrate synthase gene, Zm00001eb130080, a key enzyme responsible for synthesizing cytosolic acetyl-CoA which is subsequently used in fatty acid elongation as well as the biosynthesis of a number of metabolites including flavonoids, anthocyanins, isoprenoids, and other malonated derivatives (Fatland et al. 2002). Another strong effect locus on chromosome 3, accounting for 20% of the phenotypic variance in AUDPC, is linked to the cytochrome P450 10 (*cyp10;* Zm00001eb159250) gene. The *cyp10* gene is in the CYP72A28 family, which is important for fatty acid and isoprenoid metabolism, hormone catabolism, and cytokinin biosynthesis. The CYP72 family also encompasses the CYP716A subfamily, which is involved in triterpenoid biosynthesis (Fukushima et al. 2011). Recently, it was shown that overexpression of CYP72A28 in *Arabidopsis* altered abiotic stress response by modifying the accumulation of anthocyanins, supporting this gene’s involvement in the reddening response in the 282 diversity panel (Leopold and Thornton 2025).

Three unique loci linked to genes potentially involved in leaf reddening were identified on chromosome 5 that were associated with AUDPC and DS; one detected by BLINK for AUDPC and two by FarmCPU for AUDPC and DS. One of these SNPs was embedded in a multidrug and toxic compound extrusion 34 (*mate34;* Zm00001eb237110) gene. Although MATEs perform diverse functions, one of their most important is facilitating flavonoid transport (Pastacaldi et al. 2024). Furthermore, they are involved in mounting disease defense, including basal defense against cucumber mosaic virus (CMV) and turnip crinkle virus (Pastacaldi et al. 2024; Ishihara et al. 2008). Farther down the chromosome, a putative geranylgeranyl pyrophosphate synthase (GGPS) 4 (Zm00001eb250080) is tightly linked to an AUDPC QTN. These GGPSs catalyze the synthesis of geranylgeranyl pyrophosphate, serving as the basis for the biosynthesis of diterpenes and tetraterpenes, including carotenoids and many other plant secondary metabolites (Beck et al. 2013). The final chromosome 5 QTN is tightly linked to two genes of potential interest given their roles in pigment production. The Zm00001eb251670 gene which encodes a G2-like-transcription factor 11 (*glk11*), includes a regulation of phenylpropanoid metabolic process gene ontological (GO) term (GO:2000762). Phenylpropanoids are known to be important in floral pigment production (Deng and Lu 2017). Just a few hundred base pairs farther is an alternative oxidase 1 (*aox1*; Zm00001eb251680). The homologous gene, *ZmPTOX* on chromosome 2, regulates leaf and kernel color by modulating plastid development and may function in amyloplasts, where it could contribute to carotenoid biosynthesis (Huang et al. 2025).

Nine other significant loci were tightly linked to genes with potential roles in plant defense, at least four of which may be related specifically to virus resistance. On chromosome 1, a significant SNP associated with DS was detected by FarmCPU and tightly linked to the CCAAT-HAP5-transcription factor 54 (*ca5p4*; Zm00001eb064730) which includes an NF-YC domain. NFYs are highly conserved transcription factors consisting of NF-YA, NF-YB, and NF-YC subunits. Two of these subunits, NF-YC and NF-YA, are targets of two rice viruses, the tenuivirus, rice stripe virus, and the fijivirus, southern black streaked dwarf virus (Tan et al. 2024). Overexpression of NFYs significantly reduces the expression of genes in the jasmonic acid pathway, inhibiting antiviral defense (Tan et al. 2022). Another gene, Zm00001eb189980, encoding a protein with peptidase A1 and aspartyl protease APCB1 domains, is located just 3.3 kb from the chromosome 3 QTN. Although there are three genes with less obvious functions in response to MaYMV infection more tightly linked to the QTN, aspartyl proteases are known to play pivotal roles not only in plant virus defense, but also in fungal and bacterial defense, biotic and abiotic stress response, and plant growth. For instance, when the aspartyl protease, *OsAp77* gene was knocked out in rice, mutant plants displayed significantly higher levels of virus titer when inoculated with CMV (Alam et al. 2014).

A large effect QTN on chromosome 7 associated with DS is tightly linked to the *calreticulin 2* (*crt2*, Zm00001eb304290) gene. Calreticulins are Ca^2+^ binding proteins that localize to the endoplasmic reticulum and are involved in response to a variety of biotic and abiotic stresses (Jia et al. 2009). The importance of calreticulins in plant response to tobacco mosaic virus (TMV) has been demonstrated multiple times. In tobacco, calreticulin was found to bind the TMV movement protein *in vivo* and *in vitro*, with both proteins co-localizing to the plasmodesmata. Overexpression of calreticulin significantly impaired TMV intercellular movement, highlighting the importance of this protein for virus movement and systemic infection (Chen et al. 2005). Furthermore, silencing of two calreticulin genes, *NbCRT2* and *NbCRT3,* reduced the efficacy of N immune receptor-mediated defense and were required for the expression of induced receptor-like kinases that are needed for N-mediated hypersensitive response (HR) (Caplan et al. 2009).

BLINK also detected a chromosome 9 QTN for AUDPC. This SNP is tightly linked to the glutathione transferase 13 (*gst13*; Zm00001eb402630). GSTs play an important role in mediating the HR response upon viral inoculation. In tobacco, HR in response to TMV was shown to be preceded by lower GST activity (Fodor et al. 1997). Conversely, elevated GST activity was associated with fewer necrotic lesions and reduced levels of tobacco necrosis virus (Pogány et al. 2004), while early induction of GSTs is correlated with enhanced TMV resistance (Király et al. 2012). The importance of GSTs is highlighted in several other host-virus interactions, including *Arabidopsis* inoculated with the yellow strain of CMV (Ishihara et al. 2004), pepper inoculated with capsicum chlorosis virus (Widana Gamage et al. 2016), rice inoculated with rice yellow mottle virus or rice tungro spherical virus (Brizard et al. 2006; Satoh et al. 2013), sugar beet with beet necrotic yellow vein virus (Larson et al. 2008), as well as maize and sorghum inoculated with SCMV (Gullner et al. 1995; Wu et al. 2013).

Two moderate - strong effect QTN were identified on chromosome 4. One was tightly linked to two genes, one a multiple organellar RNA editing factor (Zm00001eb168440) and the other, Zm00001eb168430, which includes translation initiation factor eIF-2B domains. The presence of the eIF-2B domains is intriguing given their importance in plant virus resistance (Shopan et al. 2017). eIF proteins are often co-opted by viruses to complete their replication cycle (Singhal et al. 2026). In mustard greens, the eIF-2B host protein is critical for turnip mosaic virus (TuMV) protein translation. Interaction of the eIF-2Bβ subunit with a demethylation factor modifies the methylation of status of RNA encoding the TuMV viral coat protein, enhancing viral replication. However, in eIF-2B mutants the demethylation cannot occur and thus viral replication is reduced (Sha et al. 2024). The final chromosome 4 QTN is linked to several genes. The most tightly linked and characterized of which is an AT-hook (ATH) motif nuclear-localized protein (Zm00001eb204390), proteins which are involved in many important biological processes including PAMP-triggered immunity (Zhang et al. 2022).

Several QTN linked to genes with less obvious association with leaf reddening or response to virus infection were also detected. A strong effect QTN was detected by BLINK and FarmCPU for AUDPC associated with the histone h1, *hon110* gene (Zm00001eb061060) on chromosome 1. Though this gene could be involved in any number of functions, H1s have been shown to regulate defense priming and immunity in *Arabidopsis*, most likely by regulating DNA methylation and chromatin remodeling (Sheikh et al. 2023). On chromosome 2, the cytochrome b5-1 gene, *cybv1* (Zm00001eb095000) was detected by FarmCPU for AUDPC. This gene is significantly downregulated in response to drought as well as abscisic acid (Che et al. 2025), a phytohormone known for its role in abiotic stress response but is also important for plant disease resistance (Makris et al. 2026). Another minor effect QTN on chromosome 3 is associated with a bHLH-transcription factor (*bhlh179*; Zm00001eb151300) which is part of the lectin receptor-like kinase gene family which is known to mediate plant immune response (Yang et al. 2025). A final significant QTN on chromosome 10 was detected by FarmCPU for DS, but none of the genes that are most tightly linked to this locus appear to have any known function in response to virus infection, biotic stress response, or pigmentation.

No significant QTNs for ELISA were detected by any GWAS model and, in fact, none even exceeded the less stringent suggestive threshold. The QTN most closely approaching significance was located near the bottom of chromosome 7 and colocalized with a SNP exceeding the suggestive threshold for DS in the BLINK model, further supporting that this locus may have an important influence on MaYMV titer and overall DS. This QTN was most tightly linked to an E3 ubiquitin-protein ligase RKP (Zm00001eb328050), which contains a response to virus ontological term (GO:0009615). Notably, the polerovirus silencing suppressor protein, P0, functions by forming an E3 ubiquitin ligase complex with a host ubiquitin ligase and other co-factors to mark AGO1 for degradation, which is a key protein in plant RNAi-mediated antiviral immunity (Bortolamiol et al. 2007; Baumberger et al. 2007). It is unknown which maize ubiquitin ligase(s) complex(es) with MaYMV P0, but if the affinity of this interaction is decreased or lost altogether due to an alternate version of the E3 ubiquitin ligase, then silencing suppression by MaYMV would be reduced and the host RNAi machinery would be more effective in limiting MaYMV replication, thus reducing titer.

Although it may have been insufficiently sensitive for GWAS analysis of MaYMV titer in this population, the PAS-ELISA test was found to be effective at detecting infection, with 246 of the 250 lines testing positive across the two Deepot screening experiments even using a stringent cutoff threshold of 2x the healthy control absorbance. Although ELISA is semi-quantitative in nature, the overall variance in titer was compressed and ELISA failed to detect significant differences in titer between even the most MaYMV tolerant line, Ky226, and other lines including the susceptible control, Oh28, differences which were readily apparent via RT-qPCR. These results highlight the value of ELISA for disease monitoring given its high throughput economical testing but suggest that more sensitive and costly RT-qPCR assays may be a desirable choice for breeding, selection, and mapping analyses for MaYMV.

The identification of Ky226 as the most resistant line in this study is perhaps unsurprising given its high level of resistance to the related luteovirus, BYDV-PAV (Schmidt et al. 2024). The BYDV resistance gene in Ky226 was previously mapped to the bottom of chromosome 10, although the confidence interval reported by Schmidt *et al*. is nearly 3 Mbp downstream of the chromosome 10 QTN reported in this study. Interestingly, this was the only QTN for which there were no obvious candidate genes within a 50 kb interval on either side that were associated with reddening or disease responses. Whether the SNP identified in this study is associated with a MaYMV resistance gene unique from that reported for BYDV resistance previously or whether GWAS mapping resolution is limited due to insufficient diversity and/or recombination is unclear. Several adaptation and domestication related traits map to the adjacent genetic bin 10.04, which resulted in a selective sweep and strongly reduced the genetic variability in this and nearby genomic regions in modern maize (Tian et al. 2009), potentially reducing the mapping resolution in this region due to insufficient nucleotide diversity.

Although further study is required to elucidate the effects of non-symptomatic infection by MaYMV on plant vigor and overall production, this study indicates that the majority of maize lines present in the Goodman 282 diversity panel display minimal visible response to infection. Furthermore, we report at least one line, Ky226, which was non-symptomatic and had reduced titer relative to the other lines tested, that could be a valuable resource for breeding varieties with better MaYMV resistance. The identification of numerous QTN associated with genes related to the production of phenolic, carotenoid, and production of related molecules related to pigmentation elucidates the potential role of these genes in reddening symptom development, many of which have not yet been characterized. Likewise, the identification of several QTN associated with genes governing plant viral and disease defense pathways enhances our understanding of these processes and, when integrated with advanced gene-editing technologies, supports the development and engineering of corn varieties with superior disease resistance.

## Data availability

The authors affirm that all data necessary for confirming the conclusions of the article are present within the article, figures, and tables.

## Acknowledgements

The authors would like to thank Courtney McCusker (USDA-ARS) for her assistance with sample collection, Mark Jones (USDA-ARS) for his help maintaining germplasm, and Dr. Anna Whitfield (North Carolina State University) for providing the aphid colony used in this study.

## Disclaimer statement

The findings and conclusions in this publication are those of the authors and should not be construed to represent any official USDA or U.S. Government determination or policy.

---

**Fig. S1.**
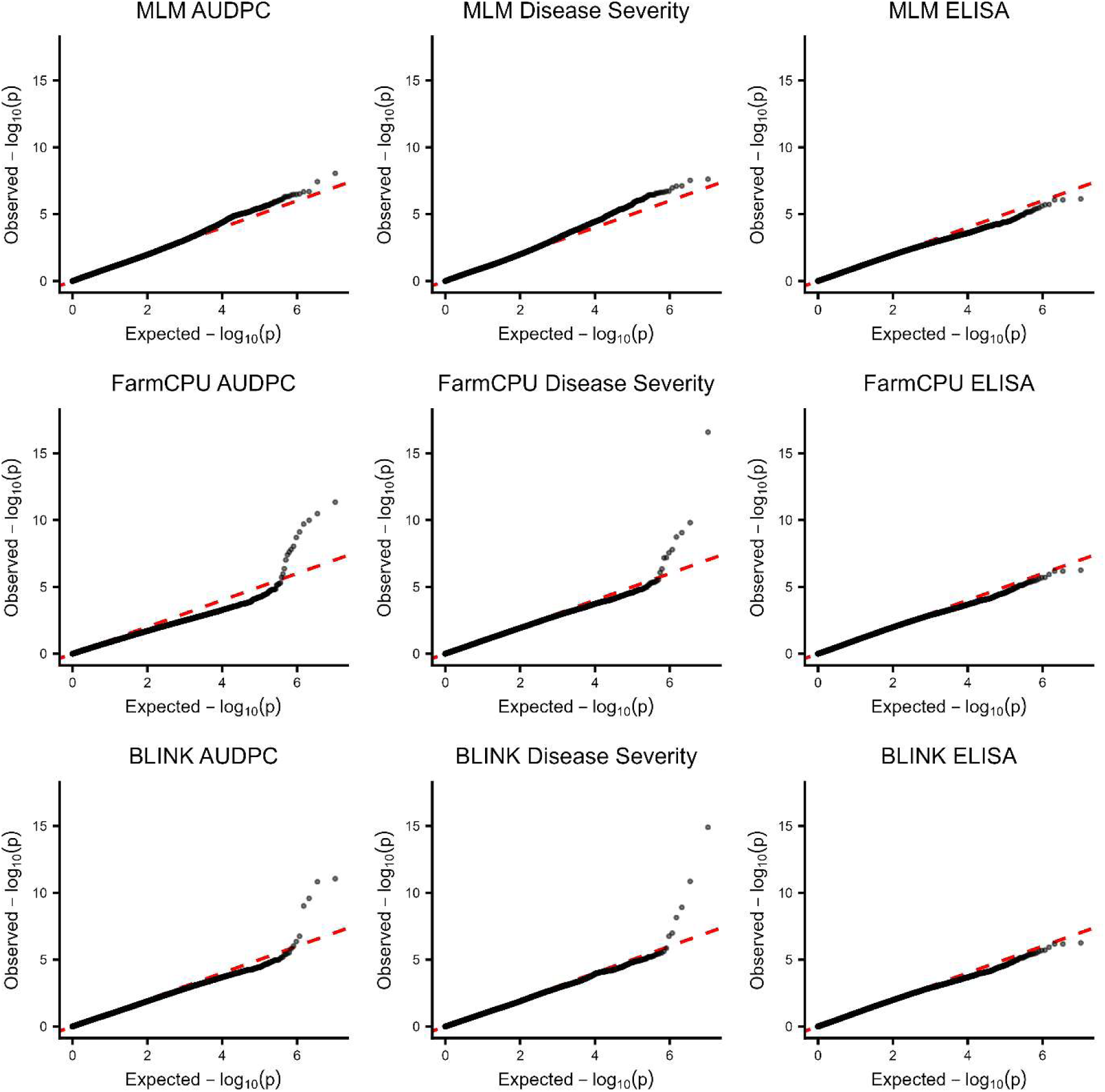
QQ-plots for response to maize yellow mosaic virus in the maize 282 association mapping panel.

